# The predictability of genomic changes underlying a recent host shift in Melissa blue butterflies

**DOI:** 10.1101/222703

**Authors:** Samridhi Chaturvedi, Lauren K. Lucas, Chris C. Nice, James A. Fordyce, Matthew L. Forister, Zachariah Gompert

## Abstract

Despite accumulating evidence that evolution can be predictable, studies quantifying the predictability of evolution remain rare. Here, we measured the predictability of genome-wide evolutionary changes associated with a recent host shift in the Melissa blue butterfly (*Lycaeides melissa*). We asked whether and to what extent genome-wide patterns of evolutionary change in nature could be predicted (1) by comparisons among instances of repeated evolution, and (2) from SNP × performance associations in a lab experiment. We delineated the genetic loci (SNPs) most strongly associated with host use in two *L. melissa* lineages that colonized alfalfa. Whereas most SNPs were strongly associated with host use in none or one of these lineages, we detected a ~two-fold excess of SNPs associated with host use in both lineages. Similarly, we found that host-associated SNPs in nature could also be partially predicted from SNP × performance (survival and weight) associations in a lab rearing experiment. But the extent of overlap, and thus degree of predictability, was somewhat reduced. Although we were able to predict (to a modest extent) the SNPs most strongly associated with host use in nature (in terms of parallelism and from the experiment), we had little to no ability to predict the direction of evolutionary change during the colonization of alfalfa. Our results show that different aspects of evolution associated with recent adaptation can be more or less predictable, and highlight how stochastic and deterministic processes interact to drive patterns of genome-wide evolutionary change.

## Introduction

Repeated evolution of similar traits in populations or species under similar ecological conditions has been widely documented, and often involves the same genes or alleles (Arendt & Reznick, 2008; Conte *et al.,* 2012; Martin & Orgogozo, 2013; Ord & Summers, 2015; Orgogozo, 2015). Such repeatability suggests patterns of evolutionary change may be predictable, either by comparison with other instances of evolution (i.e., in the context of parallel or convergent evolution) or from a mechanistic understanding of the sources and targets of selection (Lässig *et al.,* 2017). The degree to which evolution is repeatable and predictable is of general interest, as high repeatability would suggest a more central role for deterministic evolutionary processes (e.g., natural selection), and perhaps increased constraint or bias in terms of the trait combinations, developmental pathways or mutations that can result in adaptation to a given environment. Such constraints could impose general limits on patterns of biological diversity (Morris, 2008; Losos, 2011; Scotland, 2011; Speed & Arbuckle, 2017). But further progress requires moving beyond documenting instances of repeated evolution, and instead quantifying the degree to which, and context in which, evolution is repeatable or predictable, as well as identifying the factors mediating this (e.g., Soria-Carrasco *et al.,* 2014; Egan *et al.,* 2015; Erickson *et al.,* 2016; Gompert & Messina, 2016; Speed & Arbuckle, 2017).

Instances of repeated evolution provide just one of several ways to assess the predictability of evolution; the predictability of evolution can also be considered in terms of comparisons between (i) experiments linking genotype to phenotype or fitness and (ii) evolutionary patterns in natural populations (Lässig *et al.,* 2017; Agrawal, 2017). For example, field transplant experiments can be used to identify genes or traits under divergent selection between two environments, and one can then ask whether or to what extent patterns of genetic differentiation between natural populations occupying those different environments could be predicted from the experimental results (e.g., Barrett *et al.,* 2008; Soria-Carrasco *et al.,* 2014; Egan *et al.,* 2015+). This approach has received relatively little attention compared to direct tests for parallel or convergent evolution in nature (Agrawal, 2017), but it may have a greater ability to identify the mechanisms underlying predictability by better isolating components of the many evolutionary and ecological processes affecting natural populations (Losos, 2011; Gompert *et al.,* 2014a; Speed & Arbuckle, 2017). With that said, a lack of consistency between experimental and natural populations can be difficult to interpret, as experiments can miss key features of the natural environment. Whereas both of these approaches (i.e. prediction from experiments and studies of parallelism) have been used in isolation to assess the predictability of evolution, they have rarely been used in a single system and in a comparative manner (but see, e.g., Soria-Carrasco *et al.,* 2014; Egan *et al.,* 2015).

Here we consider the predictability of genome-wide evolutionary changes (hereafter genomic change) associated with a host-plant shift in the Melissa blue butterfly, *Lycaeides melissa* (Lycaenidae). We focus on a quantitative comparison of the two aspects of predictability discussed above. *Lycaeides melissa* occurs throughout western North America, where it feeds on legumes, particularly species of *Astragalus, Lupinus,* and *Glycyrrhiza. Medicago sativa* (alfalfa, a common forage crop and also a legume) was introduced to western North America in the mid 1800s, and has since been colonized by *L. melissa* (Michaud *et al.,* 1988). This is a poor host in terms of caterpillar survival, weight, and adult fecundity (Forister *et al.,* 2009, 2010; Scholl *et al.,* 2012). Nonetheless, many *L. melissa* populations persist on and have partially adapted to this plant species (e.g., on average, populations on *M. sativa* exhibit increased larval performance and adult oviposition preference relative to populations that do not feed on *M. sativa*; Forister *et al.,* 2013; Gompert *et al.,* 2015). At present, we do not know whether *L. melissa* colonized *M. sativa* once or multiple times, nor do we know whether the alfalfa-feeding populations are connected by appreciable levels of gene flow. But such information is critical for assessing the degree to which different populations or groups of populations represent independent instances of adaptation, and thus whether they can be used to quantify the repeatability of evolution.

We have additional reasons to be interested in gene flow among *L. melissa* populations. In a previous lab experiment, *L. melissa* caterpillars from populations feeding on *M. sativa* (Goose Lake Ag. = GLA; 41.9860° N, 120.2925° W) and from a population feeding on *Astragalus canadensis,* which is a native host (Silver Lake = SLA; 39.64967° N, 119.92629° W), were reared in a crossed design on either *M. sativa* or *A. canadensis* (Gompert *et al.,* 2015). We then used a multi-locus genome-wide association mapping approach to identify SNPs associated with variation in larval performance (survival and weight) for each population × host combination. This experiment showed that genetic variants associated with performance on each host plant were mostly independent, and thus, we failed to find evidence for genetic trade-offs in performance across hosts. Such trade-offs are often hypothesized to drive host plant specialization in phytophagous insects (Futuyma & Moreno, 1988; Fry, 1996). Despite the popularity of this hypothesis, very few studies have found evidence of resource-based trade-offs between hosts (but see Via & Hawthorne, 2002; Gompert & Messina, 2016). Based on these results, we raised an alternative hypothesis that host plant specialization, and particularly the loss of adaptation to an ancestral host (in this case *A. canadensis*), results from genetic drift in isolated populations that are not well connected by gene flow (similar to Grosman *et al.,* 2015). In other words, reduced performance on an ancestral host in an alfalfa feeding population could result solely from genetic drift if alleles increasing fitness on the ancestral host do not affect fitness on alfalfa, and if alfalfa feeding populations experience little to no gene flow with these ancestral populations. This hypothesis has implications for the repeatability of evolution, as it would predict a greater role for stochastic processes (i.e., genetic drift) in patterns of genomic change (i.e., evolutionary change across the genome) during repeated host shifts than would be expected if trade-offs were prevalent. Evaluating this hypothesis requires additional data on gene flow among *L. melissa* populations.

Herein, we first test whether *L. melissa* has colonized *M. sativa* one or multiple times and quantify levels of contemporary gene flow, and second quantify the predictability of genomic change associated with the colonization of *M. sativa* by *L. melissa*. Specifically, we analyze genotyping-by-sequencing (GBS) data from 26 *L. melissa* populations to ask the following questions: (i). Have *L. melissa* populations colonized the novel host *M. sativa* repeatedly in independent colonization events? (ii) To what extent do parallel genetic changes underlie repeated instances of the colonization of *M. sativa* by *L. melissa*? (iii) To what extent do SNP × larval performance associations in the aforementioned rearing experiment predict patterns of genetic differentiation between natural populations feeding on alfalfa versus native legume hosts? (iv). Is the degree of predictability higher in the context of (ii) or (iii)? See Fig. 1 for a summary of research questions and primary analyses.

**Figure 1:**
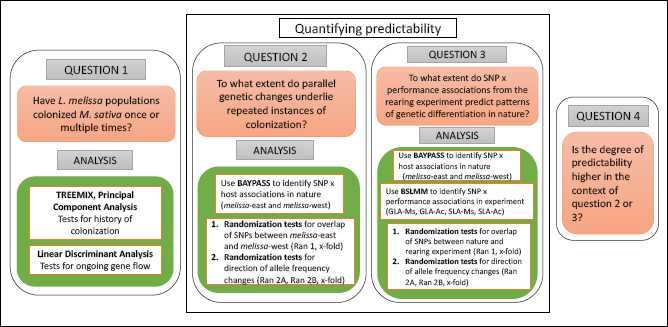
Diagram shows a schematic representation of the primary analyses conducted in this study for main objectives. Each box presents a question asked in this study and the analyses conducted to answer these questions.

## Methods

### Samples and DNA sequencing

In this study we considered GBS data from 526 *L. melissa* butterflies collected from 26 populations distributed across the western USA (Table 1). This includes 15 populations that use *M. sativa* (alfalfa) as a host, and 11 populations that use one of several native legume species (i.e., species of *Astragalus, Lupinus* or *Glycrrhiza*). GBS data from 414 of these individuals (20 populations) were previously published in a study of admixture in the *Lycaeides* species complex (Gompert *et al.,* 2014b), whereas the GBS data from the other 112 individuals (6 populations) are presented here. DNA extraction, GBS library preparation, and DNA sequencing (100 bp single-end reads with an Illumina HiSeq 2500) occurred concurrently for all 526 samples (for details refer to Gompert *et al.,* 2014a).

**Table 1:** Locality information and sample sizes for the populations included in this study. Group denotes the lineage based on `TREEMIX` results, # Ind. gives the number of individuals sequenced for this study, and Data = indicates whether the sequence data were included in Gompert *et al.* (2014b) = “2014”, or are being presented here for the first time = “Present”.

|  | Locality | Host-plant | Long. (W) | Lat. (N) | Group | # Ind. | Data |
| --- | --- | --- | --- | --- | --- | --- | --- |
| 1 | Bishop, CA (BHP) | *Glycyrrhiza sp.* | 118.28 | 37.17 | *melissa-west* | 20 | 2014 |
| 2 | Crystal Creek Park, NV (VCP) | *Medicago sativa* | 119.99 | 39.51 | *melissa-west* | 20 | Present |
| 3 | Gardnerville, NV (GVL) | *Medicago sativa* | 119.78 | 38.81 | *melissa-west* | 18 | 2014 |
| 4 | Red Earth Way, NV (REW) | *Medicago sativa* | 118.84 | 38.98 | *melissa-west* | 20 | 2014 |
| 5 | Silver Lake, NV (SLA) | *Astragalus canadensis* | 119.93 | 39.66 | *melissa-west* | 18 | 2014 |
| 6 | Sierra Valley, CA (SVY) | *Medicago sativa* | 121.14 | 39.09 | *melissa-west* | 20 | 2014 |
| 7 | Trout Pond Trailhead, CA (TPT) | *Lupinus sp.* | 116.58 | 32.97 | *melissa-west* | 13 | Present |
| 8 | Washoe Lake, NV (WLA) | *Astragalus candensis* | 118.82 | 38.65 | *melissa-west* | 20 | 2014 |
| 9 | Abel Creek, NV (ABC) | *Lupinus sp.* | 117.65 | 41.44 | *melissa-east* | 19 | Present |
| 10 | Brandon, SD (BSD) | *Medicago sativa* | 96.54 | 43.63 | *melissa-east* | 20 | Present |
| 11 | Cody, WY (CDY) | *Medicago sativa* | 108.98 | 44.51 | *melissa-east* | 23 | 2014 |
| 12 | Cokeville, WY (CKV) | *Medicago sativa* | 110.90 | 42.01 | *melissa-east* | 10 | 2014 |
| 13 | De Beque, CO (DBQ) | *Medicago sativa* | 108.21 | 39.32 | *melissa-east* | 20 | 2014 |
| 14 | Deeth-Charleston, NV (DCR) | *Lupinus sp.* | 115.38 | 41.30 | *melissa-east* | 20 | 2014 |
| 15 | Goose Lake, CA (GLA) | *Medicago sativa* | 120.29 | 41.30 | *melissa-east* | 20 | 2014 |
| 16 | Lander, WY (LAN) | *Medicago sativa* | 108.36 | 42.65 | *melissa-east* | 24 | 2014 |
| 17 | Lamoille Canyon, NV (LCN) | *Lupinus sp.* | 115.47 | 40.68 | *melissa-east* | 20 | 2014 |
| 18 | Montrose, CO (MON) | *Medicago sativa* | 107.82 | 38.37 | *melissa-east* | 20 | 2014 |
| 19 | Montague, CA (MTU) | *Medicago sativa* | 122.53 | 41.73 | *melissa-east* | 19 | 2014 |
| 20 | Ophir City, NV (OPC) | *Lupinus sp.* | 117.24 | 38.94 | *melissa-east* | 19 | 2014 |
| 21 | Star Creek Canyon, NV (SCC) | *Lupinus sp.* | 118.12 | 40.55 | *melissa-east* | 16 | 2014 |
| 22 | Surprise Valley, CA (SUV) | *Medicago sativa* | 120.10 | 41.28 | *melissa-east* | 20 | 2014 |
| 23 | Upper Alkali Lake, CA (UAL) | *Medicago sativa* | 120.15 | 41.74 | *melissa-east* | 20 | Present |
| 24 | Victor, ID (VIC) | *Medicago sativa* | 111.11 | 43.66 | *melissa-east* | 20 | 2014 |
| 25 | Yellow Pine, WY (YWP) | Unknown native legume | 105.40 | 41.25 | *melissa-east* | 20 | 2014 |
| 26 | Albion Meadows, UT (ABM) | *Lupinus sp.* | 111.92 | 40.48 | *melissa-east* | 46 | 2014 |

### Genome alignment and genetic variation in populations

We used the `aln` and `samse` algorithms from `bwa` 0.7.5a-r405 to align 100-bp single-end reads (525 million reads) to our draft *L. melissa* genome (the draft genome is described in Gompert *et al*., 2015). This included re-aligning the data from Gompert *et al.* (2014b) as those results preceded the current genome assembly and annotation. We allowed a maximum of four differences between each sequence and the reference (no more than two differences were allowed in the first 20 bp of the sequence). We trimmed all bases with a phred-scaled quality score lower than 10 and only placed sequences with a unique best match in our data. We then used `samtools` (version 0.1.19) to compress, sort and index the alignments (Li et *al.,* 2009). We identified (verified) single nucleotide variants and calculated genotype likelihoods, but considering only the set of SNPs identified previously by Gompert *et al.*(2015). The original variant set was called using many of the L. *melissa* samples included here, as well as butterflies from the rearing experiment described above. By focusing on this variant set, we ensured that clear comparisons could be made between the data from the experimental and natural populations in terms of tests of predictability. Variants were called using `samtools` and `bcftools` (version 0.1.19) and were only output if the posterior probability that the nucleotide was invariant was less than 0.01 (with a full prior with *θ* = 0.001), and if data were present for at least 80% of the individuals. All SNPs from Gompert *et al.* (2015) were also identified as SNPs in the current data set based on these criteria, resulting in 206,028 high-quality SNPs. These SNPs had an average sequencing depth of 17.29 (SD = 11.0) per individual. We used an expectation-maximization algorithm to obtain maximum-likelihood estimates of population allele frequencies while accounting for uncertainty in genotypes (based on the calculated genotype likelihoods from `bcftools`; Li, 2011; Soria-Carrasco *et al.,* 2014).

### Colonization history and tests for gene flow

We used a series of analyses to assess the degree of independence in evolutionary change across the *M. sativa* feeding populations. We were interested in independence both in terms of historical colonization and admixture/gene flow, and in terms of contemporary gene flow. We first used principal components analysis (PCA) as an ordination-based approach to examine whether the *M. sativa* populations formed a single coherent cluster in genotype space, as would be predicted if alfalfa was colonized a single time. We ran the PCA in `R` (version 3.4.1) on the among individual similarity (i.e., genetic covariance) matrix, which was calculated from genotype point estimates for 14,051 common (global minor allele frequency > 5%) SNPs (genotypes were inferred using a mixture model for allele frequencies with k = 2 to 5 source populations, as in Gompert *et al.,* 2014b). We then used `TREEMIX` (version 1.12) to construct a population graph depicting the relationships among the focal populations (Pickrell & Pritchard, 2012). This method first fits a bifurcating tree based on the population allele frequency covariance matrix (based on the maximum likelihood allele frequency estimates), and then adds migration/admixture edges to the tree to improve the fit. Thus, it allowed us to test for both the monophyly of *M. sativa*-feeding populations (i.e., to test whether there were one or multiple successful colonization events), and to ask whether, if alfalfa was colonized multiple times, the populations have since experienced appreciable historical gene flow/admixture which would reduce their evolutionary independence. We rooted the population tree with two *Lycaeides anna* populations, which were set as the outgroup (data from Gompert *et al.,* 2014b) and fit graphs allowing 0-10 admixture events. We calculated the proportion of variance in allele frequency covariances explained by the population graph with varying numbers of admixture events to quantify model fit, and to determine whether individual admixture events substantially improved model fit (Pickrell & Pritchard, 2012).

We then used stochastic character mapping to estimate the number of host shifts to *M. sativa* based on the tree from `TREEMIX` (Bollback, 2006). We treated host use (native host vs. *M. sativa*) as a trait for ancestral character state reconstruction (as in, e.g., Dobler *et al.,* 1996; Hwang & Weirauch, 2012). We fixed the root of the tree as native feeding because of the known recent introduction of *M. sativa* to North America. We used the `make.simmap` function in the R package phytools (version 0.6-44) for this analysis (Revell, 2012), and based our inference on two Markov chain Monte Carlo (MCMC) runs each with a 10,000 iteration burn-in, 100,000 sampling steps, and a thinning interval of 50. The probabilistic character state simulations used to estimate the number of shifts to *M. sativa* incorporated uncertainty in the character transition matrix.

We then used an assignment-based approach, namely discriminant analysis, to identify individuals that were likely migrants from another population. Our goal here was to assess evidence of contemporary gene flow in terms of actual migrants (we were not attempting to detect later generation hybrids or estimate admixture proportions; the latter can be found in Gompert *et al.,* 2014b). We used the `lda` function from the `MASS` package in `R` to assign individuals to populations based on the first four PCs of the genotypic data (see above; these accounted 95% of the genetic variation), and this was done in a pair-wise manner for all populations, although we were most interested in pairs of adjacent populations. We used k-fold cross-validation to estimate assignment probabilities. Results from this set of analyses (described in detail in the Results below) indicated that *L. melissa* have colonized *M. sativa* at least twice (and probably more times than that), once in the western Great Basin and once in the central/eastern Great Basin and Rocky Mountains, and that there has been little gene flow between these groups of populations (see e.g., Fig. 2). We thus use these two groups of populations, hereafter referred to as *melissa*-west and *melissa*-east, respectively, to quantify the extent of parallel genomic change associated with alfalfa-colonization and adaptation (experimental evidence of adaptation to alfalfa in general comes from, e.g., Forister *et al.,* 2013; Gompert *et al.,* 2015).

**Figure 2:**
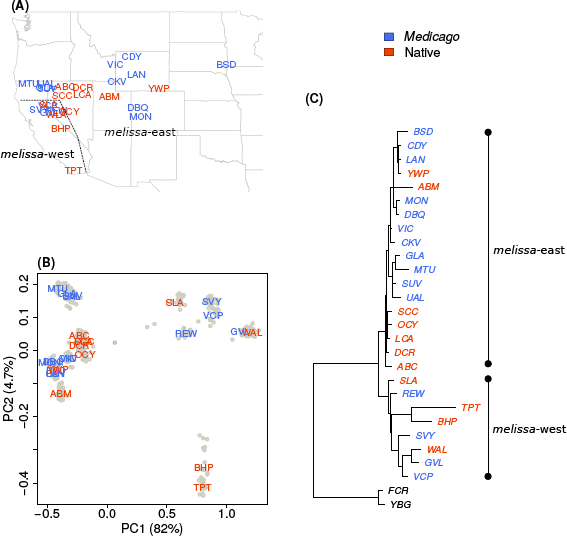
(A) Map shows sample locations with populations colored based on host association. Population labels correspond to abbreviations for geographical locations in Table 1, and the line separates populations belonging to the eastern and western clades. (B) Plot shows summary of population structure based on principal component analysis. Abbreviations indicate populations corresponding to the map (A). The points denote individuals in each population used for the analysis. (C) Population graph from `TREEMIX` for *L. melissa* populations used in this study (N=26), allowing one migration or admixture event (the actual migration edge from the outgroup to ABM is not shown). Terminal nodes are labeled by abbreviations for geographical locations from where samples were collected and colored according to host-plant association.

### Quantifying the predictability of genomic change

We measured and compared the predictability of genomic change associated with colonization of alfalfa by *L. melissa* in two ways: (i) the degree of parallelism in genomic change during two independent host shifts onto *M. sativa* in nature, and (ii) how well patterns of genomic change in nature could be predicted from performance × SNP associations in a rearing experiment. We did this by testing for and quantifying an excess overlap in SNPs associated with host use, that is, the SNPs with the greatest allele frequency differences between native and alfalfa-feeding populations in nature and the SNPs most strongly associated with performance in the rearing experiment. We report these values as x-fold enrichments. As an example, an x-fold enrichment of 2.0 would imply that twice as many SNPs are associated with host use in, e.g., repeated instances of colonization of *L. melissa,* as expected by chance (see details of null models below) and thus would mean that exceptional patterns of genomic change can be predicted from one colonization event to the other twice as well as would be the case with no information. We considered x-fold enrichments as measures of predictability both in terms of the SNPs showing host association and in terms of the direction of these effects. In other words, we distinguished between being able to predict host-associated SNPs, and being able to predict the direction of the association. As populations will necessarily vary in the details of linkage disequilibrium (LD) between causal variants and genetic markers (such as in our SNP set), the former might be more predictable than the later (see the Discussion for details).

### Delineating SNP × host use associations in nature

We first delineated the SNPs most strongly associated with feeding on *M. sativa* in nature. We used the software package `BAYPASS version` 2.1 (Gautier, 2015) to do this by identifying SNPs with the greatest allele frequency differences between populations feeding on *M. sativa* and those feeding on native hosts; this method controls for background population genetic structure. We were interested in these SNPs as they presumably exhibited the greatest change in allele frequencies following the colonization of alfalfa, and some subset of them might be in LD with causal variants affecting host (alfalfa) adaptation (given the sparsity of GBS data, we doubt that any of these SNPs directly confer host adaptation, but this is not critical for our questions and approach).

The `BAYPASS` software used here is based on the `BAYENV` method introduced by Günther & Coop (2013). `BAYPASS` uses a hierarchical Bayesian model with a binary auxiliary variable to classify each locus (i.e., SNP) as associated or unassociated with some environmental covariate. The model attempts to control for population genetic structure by approximating the history of the populations with an allele frequency variance-covariance matrix. We ran `BAYPASS` (Gautier, 2015) with three sets of populations: (i) all populations, (ii) 8 *melissa*-west populations, and (iii) 17 *melissa*-east populations. We treated host use, coded as a binary variable indicating whether or not a population was on alfalfa, as the environmental covariate, and ran the standard covariate model. For each data set, we ran four MCMC simulations, each with a 20,000 iteration burnin and 50,000 sampling iterations with a thinning interval of 100. The regression coefficient (*β_i_*) describing the association of each SNP (*i*) with host use (after controlling for population structure) was calculated using the default option of importance sampling, which also allows for computation of Bayes factors. Bayes factors were used to compare the marginal likelihoods of models with non-zero versus zero values of *β_i_*.

To further characterize the top host-associated SNPs, we conducted additional tests wherein we asked whether the SNPs most associated with host use were overrepresented on the Z chromosome (in butterflies males are ZZ and females are ZW, and the Z chromosome tends to harbor an excess of QTL for adaptive traits; Sperling, 1994; Janz, 2003), or whether they were enriched for specific gene ontology (GO) classifications. Such enrichments might be expected if the top host-associated SNPs were indeed tagging (via LD) genetic regions affecting host use. We defined “host-associated SNPs” for these and subsequent tests as those with the largest Bayes factors from the `BAYPASS` analysis. We did this using empirical quantiles, and considered a range of cut-offs, from the top 0.1% to the top 0.01% of SNPs (with increments of 0.01%). Considering multiple quantile cut-offs (here and in additional analyses described below) let us evaluate the sensitivity of our results to particular empirical quantiles. We used a new linkage map (Gompert *et al.,* manuscript in prep.) to classify SNPs as autosomal, Z-linked, or unknown. SNPs were classified as in coding regions (exons only), genic (in gene exons or introns), or intergenic based on the structural annotation described in Gompert et *al.* (2015). GO annotation were based on 14713 `PFAM-A` matches from `INTERPROSCAN`; GO terms were assigned to SNPs within 1 kb of annotated genes. Randomization tests were used to quantify and assess the significance of enrichments for each quantile cut-off and all three data sets, all 25 populations, *melissa*-east, and *melissa*-west, and in each case 1000 randomizations were conducted.

### Tests of parallel genomic change in nature

Our first framework for quantifying predictability was to test for parallel evolution of host use between two groups of *L. melissa* populations. Following the `TREEMIX` results, we used the *melissa-east* (N=17) and *melissa-west* (N=8) populations to ask if parallel genetic changes underlie host plant use in these butterfly populations. We used randomization tests (10000 randomizations per test) to generate null expectations for the proportion of top host-associated SNPs shared between melissa-east and *melissa*-west populations, and tested if this was more than expected by chance (x-fold enrichments). Herein, we refer to this procedure as *ran1* (see Fig. 1). We performed this randomization procedure twice, first with raw Bayes factors and again using residuals from regressing Bayes factors on mean allele frequencies (averaged over the relevant populations) (we focus on the latter in the Results). We repeated *ran1* considering the top 0.01% to 0.1% (with 0.01% increments) host-associated SNPs to determine whether the degree of parallelism (i.e., predictability in the context of repeated evolution) was robust to different cut-offs for defining host-associated SNPs.

Next, we asked whether the top host-associated SNPs that were shared between *melissa*-east and *melissa*-west populations showed a consistent direction in terms of the allele frequency difference between populations feeding on alfalfa versus those on native hosts. We would expect differences in a consistent direction if the same allele was favored in both colonization events and if patterns of LD (including the sign of D, the coefficient of linkage disequilibrium) between causal variants and our SNP markers were consistent between these groups of populations. We tested for consistency in the sign of allele frequency differences using both raw allele frequencies and standardized allele frequencies from `BAYPASS`; the latter are residuals after controlling for background population structure (we focus on the latter in the Results). For each SNP and group *(melissa*-east or *melissa*-west) we calculated 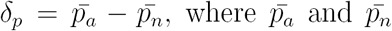 are the mean (raw or standardized) allele frequencies for the alfalfa and native-feeding populations, respectively. For the top shared host-associated SNPs, we enumerated the cases where the sign of the allele frequency difference *(δ_p_)* was the same in *melissa*-east and *melissa*-west. We conducted two sets of randomization tests to compare this to null expectations. First, we asked whether the number of shared top host-associated SNPs with the same sign for *δ_p_* was greater than expected if the *δ_p_* vectors in *melissa*-east and *melissa*-west were independent. We did this by permuting one of the sign vectors; we refer to this procedure as *ran2A* (see Fig. 1). In an additional randomization test (hereafter *ran2B*), we asked whether a greater proportion of the shared top host-associated SNPs had the same sign for *5_p_* than expected based on sign overlap for the rest of the SNPs. This was done by permuting the classification of SNPs as shared top host-associated or not.

### Predictability of patterns in natural populations from experimental outcomes

We next asked how well SNP × host association in nature can be predicted from SNP × performance association from a published lab experiment (Gompert *et al.,* 2015). In (Gompert *et al.,* 2015), we quantified the association between each SNP and host-specific survival or adult weight as a model-averaged locus effect (MAE) by fitting Bayesian Sparse Linear Mixed Models (BSLMMs). This method includes a genetic relatedness matrix and the genotype of each individual at each SNP as predictors of each individual’s phenotype (Zhou *et al.,* 2013). MAEs are given by the formula 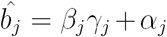, where *β_j_* is SNP *j*’s main effect if it is included in the model (i.e., if it has a main effect), *ϒ_j_* is the posterior inclusion probability for SNP *j* (i.e. the probability that SNP *j* has a main effect, that is, that it tags a causal variant), and *α_j_* is SNP *j*’s contribution to the phenotypic variation via the genetic relatedness matrix. In this experiment, we estimated MAE for the following treatments: (i) larvae from GLA (which feeds on *M. sativa*) reared on *M. sativa* (GLA-Ms) 2) larvae from GLA reared on *A. canadensis* (GLA-Ac) 3) larvae from SLA (which feeds on *A. canadensis*) reared on *M. sativa* (SLA-Ms) and 4) larvae from SLA reared on *A. canadensis* (SLA-Ac). We used these MAEs to delimit SNPs with the greatest association with host-specific performance in the experiment. We then asked whether these SNPs also showed substantial allele frequency differences between alfalfa and non-alfalfa feeding populations in nature. We enumerated the SNPs that were top host-associated SNPs in the BAYPASS analysis and that were top performance-associated SNPs (based on the MAEs). We considered classifications based on the top 0.1% to 0.01% of SNPs and based on each experimental treatment and performance metric (weight or survival) (different SNPs were associated with performance on each host; Gompert *et al.,* 2015). Comparisons were made between the experimental results and `BAYPASS` results for: *melissa*-east, *melissa*-west and all populations. We used the *ran1* approach (Fig. 1) to test for and quantify an excess of overlap between the top host-associated (in nature) and top performance-associated (in the experiment) SNPs.

Next, we asked whether, for the shared top host-associated and top performance-associated SNPs, the direction of allele frequency differences in nature *(δ_p_*, see the previous section) was consistent with the direction of the SNP × performance association from the experiment (see Fig. S1). For example, an allele associated with increased survival on alfalfa in the experiment would be predicted to be at higher frequency in the alfalfa-feeding populations. We considered all combinations of definitions for top host-associated SNPs (melissa-east, *melissa-west,* all populations) and top performance-associated SNPs (all experimental treatments and both weight and survival), and in each case enumerated the instances where the sign for *δ_p_* and the sign of the MAE were consistent. Then, as we did for the tests of parallelism in nature (see preceding section), we used randomization tests to ask whether and to what extent there was more consistency than expected (i) assuming the direction of SNP × host and SNP × performance association were independent *(ran2A),* and (ii) based on the consistency of *δ_p_* and MAEs for all other SNPs *(ran2B*).

Even if SNP × performance associations were not generally predictive of SNP × host association in nature, they could be locally predictive of exceptional genetic differentiation between the specific populations used in the rearing experiment. To test this, we first quantified genetic differentiation between GLA and SLA at each SNP locus using Hudson’s estimator of F_ST_ (Bhatia *et al.,* 2013). Then, similar to the analyses described above, we identified the most differentiated loci between GLA and SLA, that is the 0.1% to 0.01% most differentiated SNPs, and tested for significant overlap between these SNPs and the top performance-associated SNPs *(ran1)*, and for whether the direction of allele frequency difference between these populations was consistent with the direction of the SNP × performance association from the experiment *(ran2A* and *ran2B)*. We then repeated these analyses with the most differentiated SNPs between GLA and a relatively close native-feeding population (ABC, host = *Lupinus,* distance from GLA = 220.8 km), and with SLA and a nearby alfalfa-feeding population (VCP, distance from SLA = 17.5 km) (GLA and SLA are themselves 184.9 km apart). Here, we considered only the performance-associated SNPs from GLA (for the GLA × ABC comparison) or the performance-associated SNPs from SLA (for the SLA × VCP comparison).

## Results

### Colonization history and tests for gene flow

Ordination with PCA indicated that most (86.7%) of the genetic variation in the samples was accounted for by the first two principal components. These PCs largely separated individuals and populations based on geography rather than host plant (Fig. 2). The best bifurcating tree from `TREEMIX` explained 94.3% of the variance in population covariances. Consistent with the PCA results, *L. melissa* populations formed two major clades that grouped populations by geography; each major clade included a mixture of populations feeding on alfalfa and native hosts (Fig. 2C). Adding migration edges to the tree increased the variance explained (Fig. S2), with the biggest gain from a single migration edge. This tree (graph) explained 97% of the variation in the data and allowed for gene flow from the outgroup *Lycaeides anna* to a single high-elevation *L. melissa* population at Albion meadows, UT. As such gene flow is unlikely in terms of geography, and because this population is phenotypically distinct from other *L. melissa*, it was excluded from further analyses. Stochastic character mapping of host use on the `TREEMIX` tree suggested four shifts from native hosts to *M. sativa* (95% credible intervals [CIs] = 2-10, posterior probability of two or more shifts = 0.95), with seven likely reversals back to native feeding (95% CIs = 4-11) (Fig. S3). Results from these analyses indicate that *L. melissa* have colonized *M. sativa* at least twice (and probably more times than that), once in the western Great Basin and once in the central/eastern Great Basin and Rocky Mountains, and that there has been little historical gene flow between these groups of populations.

With discriminant analysis, most individuals were confidently assigned to the population from which they were sampled (average assignment probability to the population of origin = 0.9918; Figs. S4, S5). Mean assignment probabilities to the population of origin were similar for same (0.964, sd = 0.0524) and different (0.984, sd = 0.953) host comparisons. Very few individuals were confidently assigned to the alternative population, that is, the one they were not sampled from (0 in 221 population pairs, 1 in 66 pairs, and 2 in 13 pairs, assignment prob. > 0.9), and we never had more than two individuals assigned to the population they were not collected from (Fig. S5). Based on all of these results we used the two clades which included populations located in the eastern and western geographical ranges of the species (hereafter *melissa-east* [N=17 populations] and melissa-west [N=8 populations]) to test for predictable genetic changes underlying host plant use (Fig. 2A, Table 1).

### Delineating SNP × host use associations in nature

Before formally quantifying the predictability of genomic change, we identified SNPs associated with host plant use in *L. melissa* populations, which we refer to as “host-associated SNPs”. Most SNPs across *melissa*-east and *melissa*-west populations had low Bayes factors (Fig. S6). However, Bayes factors were large (i.e., > 5, meaning the likelihood of the host-association model was at least five times greater than the null model) for some SNPs (1068 in *melissa*-west and 1611 in *melissa*-east) (Fig. S6).

Here we report the results for top 0.01% host-associated SNPs (N=2061). For all populations, an excess of host-associated SNPs were present on the Z-chromosome (obs. = 195, x-fold enrichment = 2.26, *P <* 0.01; randomization test). Similar results were seen for *melissa-east* (obs. = 193; x-fold enrichment = 2.23, *P* < 0.01; randomization test) and *melissa-west* populations (obs. = 134; x-fold enrichment = 1.55, *P* < 0.01; randomization test) (also see Table S1). A significant excess of the host-associated SNPs were also present in gene exons (for all populations; x-fold = 1.45; *P* < 0.01; randomization test, Tables S2, S3). These results held for melissa-east (x-fold enrichment = 1.46; P < 0.01; randomization test) and *melissa*-west (x-fold enrichment = 1.46; P < 0.01; randomization test) populations. Gene ontology (GO) analysis revealed that the host-associated SNPs are present in regions of the genome containing genes involved in a range of biological and cellular processes, and exhibit various molecular functions (Tables S4, S5, S6). Some GO terms are over-represented among the top host-associated SNPs, but not enough so to warrant particular attention at this time.

### Parallel genomic change in nature

For the top 0.01% SNPs (N=2061) with the largest Bayes factors in *melissa*-east or *melissa*-west from `BAYPASS` analysis, there was a significant excess of overlap, such that more SNPs were highly associated with host use in melissa-east and *melissa*-west than expected by chance *(ran1,* obs. = 58 shared SNPs, expected = 21, x-fold enrichment = 2.82, *P <* 0.01; Fig. 3). Six of the 58 shared top host-associated SNPs were on the Z-chromosome, which is also an excess relative to null expectations (randomization test, x-fold = 2.43; P = 0.03; Table S1). Nonetheless, the majority of top host-associated SNPs differed between *melissa*-east and *melissa*-west, resulting in a low overall correlation in Bayes factors (Pearson r = 0.06; P < 0.01). We found that the x-fold enrichments for shared top host-associated SNPs held across a range of empirical quantiles, with the greatest excess seen in the most extreme quantiles (Figure 4).

**Figure 3:**
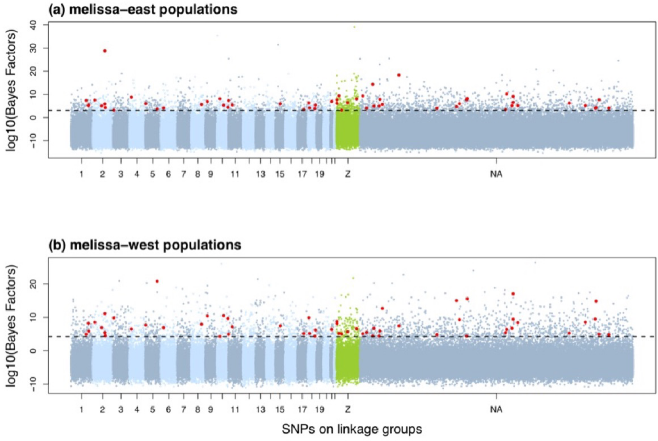
Manhattan plot shows SNPs from (a) *melissa-east* (N=17) and (b) melissa-west (N=8) population groups, along linkage groups. The horizontal dashed line delineates the top 0.01% SNPs with the highest Bayes factors. Red points denote the 58 SNPs shared by the two groups. NA indicates SNPs which did not map on any linkage group.

**Figure 4:**
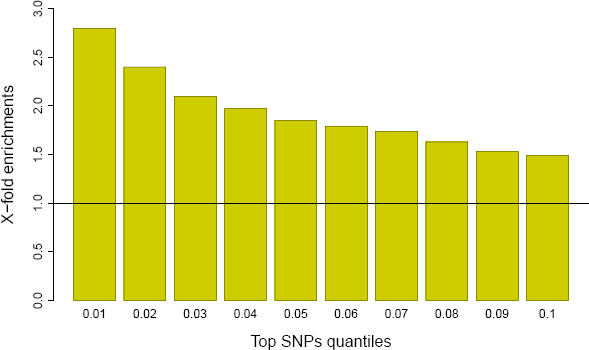
Figure 4: Barplot shows x-fold enrichments for shared SNPs between *melissa-east* and *melissa-west* populations. Results are shown for different quantile cut-offs for defining the top host-associated SNPs. The null expectation is shown with a solid horizontal line.

We found minimal evidence of concordance in the direction of allele frequency differences between alfalfa and native-feeding population when comparing *melissa*-east and *melissa*-west and considering the shared top host-associated SNPs. Specifically, for the top 0.01% SNPs (N=2061) with the largest Bayes factors in both population groups, we found no evidence of greater than expected concordance in the sign of allele frequency differences between alfalfa and native-feeding populations *(ran2A)* based on standardized or raw allele frequencies (standardized: x-fold = 1.04, P = 0.222; raw: x-fold = 1.05, P = 0.257). For the same empirical quantile, we found limited and weak evidence of greater sign coincidence for the shared top host-associated SNPs than random SNPs *(ran2B*) based on the standardized allele frequencies (x-fold = 1.05, P = 0.05), and a slight excess of sign coincidence based on the raw allele frequencies (x-fold = 1.21, P = 0.046).

### Predictability of natural processes from experimental outcomes

We found some cases where there was greater overlap than expected by chance between SNPs most associated with performance in the rearing experiment (top performance-associated SNPs) and those most associated with host use in nature (top host-associated SNPs), but this depended on the specific comparison being considered (here we again focus on results for top 0.01% SNPs, N=2061, but also provide results for other quantiles graphically; Figs. 5, 6, S7). For example, we found an excess overlap in top survival-associated SNPs for GLA reared on *M. sativa* and top host-associated SNPs for all *L. melissa (ran1,* obs. = 14, x-fold enrichment = 1.74X, P = 0.03), but not for *melissa*-east or *melissa*-west (these were marginally significant with P = 0.06 and 0.07, respectively). We found excess overlap in top survival-associated SNPs for GLA reared on *A. canadensis* and top host-associated SNPs for (i) *melissa-*east (*ran1,* obs. = 16, x-fold enrichment = 1.95X, P = 0.01), and (ii) *melissa-west (ran1,* obs. = 14, x-fold enrichment = 1.71X, P = 0.04) (Table S7). Survival-associated SNPs in SLA were not predictive of host-associated SNPs in nature, but we did detect an excess of shared weight-associated SNPs in SLA and host-associated SNPs. For example, top weight-associated SNPs for SLA when reared on *A. canadensis* overlapped more than expected by chance with melissa-east host-associated SNPs *(ran1,* obs. = 17, x-fold enrichment = 2.23; *P* = < 0.01; Table S8). In addition, for top weight-associated SNPs for SLA reared on *M. sativa* overlapped more than expected by chance with *melissa*-west host-associated SNPs *(ran1,* obs. = 19, x-fold enrichment = 2.5; P < 0.01; Table S8). We found a single case of excess overlap in top weight-associated SNPs for GLA reared on *M. sativa,* which was with the top host-associated SNPs for *melissa*-west *(ran1,* obs. = 16, x-fold enrichment = 2.06X, *P* = 0.01).

**Figure 5:**
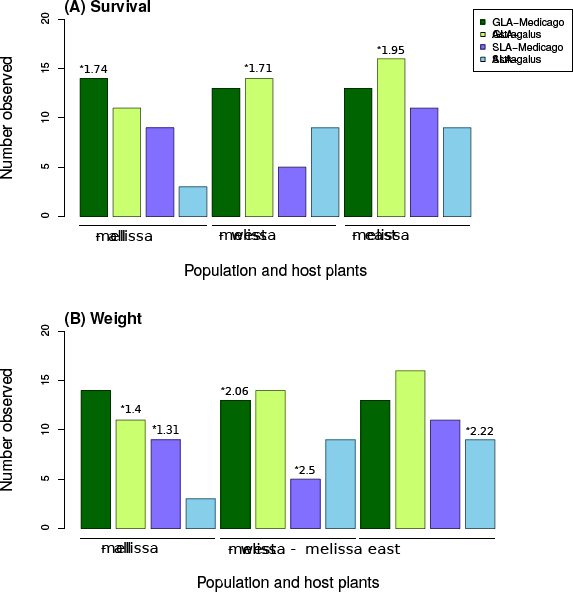
Barplots show observed number of overlapping SNPs between performance-associated SNPs in the rearing experiment and host-associated SNPs (x-axis) in nature for the top 0.01% empirical quantile. In the figure legend, GLA-Medicago indicates larvae from GLA reared on *M. sativa,* GLA-Astragalus indicates larvae from GLA reared on *A. canadensis,* SLA-Medicago indicates larvae from SLA reared on *M. sativa,* and SLA-Astragalus indicates larvae from SLA reared on *A. canadensis.* * indicates x-fold enrichments with *P ≤* 0.05.

**Figure 6:**
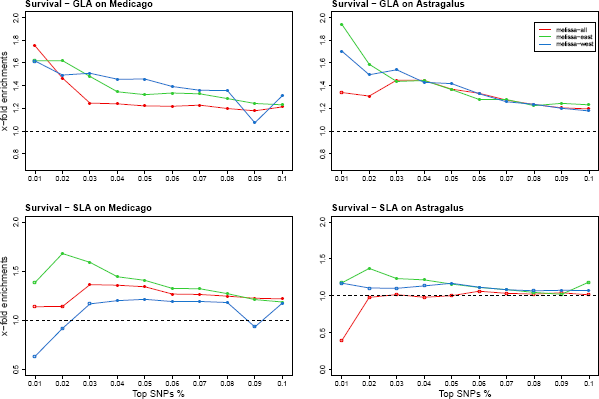
Line plots show x-fold enrichments across quantiles for overlapping SNPs between survival-associated SNPs in the rearing experiment and host-associated SNPs in nature. Open circles indicate P > 0.05 and filled circles indicate P < 0.05.

For the top 0.01% SNPs (N=2061) host-associated SNPs and performance-associated SNPs, we found weaker evidence for and a lesser degree of concordance in the direction of allele frequency differences between alfalfa and native-feeding populations (i_p_) and signs of model average effects (MAE) of performance-associated SNPs (Figs. S8, S9, S10, S11; Tables S9, S10). Most notably, there was a modest excess of sign coincidence for performance (survival)-associated SNPs for SLA reared on *M. sativa* and host-associated SNPs in *L. melissa* east *(ran2A* and *ran2B,* x-fold enrichment = 1.39-1.59X, P ≤ 0.01) and west *(ran2A* and *ran2B,* x-fold enrichment = 1.42-1.53X, P ≤ 0.03). We also found excess sign concordance for survival-associated SNPs for GLA reared on *M. sativa* and host associated SNPs in *L. melissa* east *(ran2A,* x-fold enrichment = 1.21, P = 0.05; *ran2B,* x-fold enrichment = 1.38, P = 0.04; Table S9).

We generally found greater overlap and sign-consistency between performance-associated SNPs and the most differentiated SNPs between GLA and SLA in nature than between performance-associated SNPs and more broadly host-associated SNPs in nature (described in the previous two paragraphs). We found substantial overlap between the top 0.01% of performance-associated and the top 0.01% most differentiated SNPs (F_ST_ ≥ 0.23), with x-fold enrichments ranging from 4.53 (weight-associated SNPs for SLA reared on *A. canadensis*) to 1.42 (survival-associated SNPs for SLA reared on *A. canadensis*), with P < 0.05 for all but one of these comparisons (Table S11). Somewhat weaker overlap was detected when considering the most differentiated SNPs between GLA and ABC or SLA and VCP (x-fold enrichment = 0.77-2.06), but still in most cases the overlap was greater than expected by chance. And in general, the results were consistent across different top-SNP quantiles (Fig. S12). Results for tests of sign coincidence were more idiosyncratic, but with most cases of significant excess overlap between the sign of genetic differentiation between populations and the sign of the SNP × performance effect estimate involving experimental populations reared on *M. sativa*, though there was also some evidence of significant excess involving survival-associated SNPs for SLA reared on *A. canadensis* (Tables S12). With that said, x-fold enrichment across all comparisons never exceeded 1.55X (for *ran2B,* F_ST_ for GLA vs. SLA × weight-associated SNPs for GLA reared on *M. sativa*). Similar results were obtained for other quantile cut-offs (Figs. S13, S14).

## Discussion

Several studies have shown that evolution can be predicted, at least in part, but fewer studies have quantified the degree of predictability, and in general less attention has been paid to the predictability of genomic changes underlying complex life history traits (Speed & Arbuckle, 2017). Here we first showed that *L. melissa* butterflies have colonized a novel host plant (alfalfa, *M. sativa*) two or more times, with little to no gene flow connecting the two clades of alfalfa-feeding butterflies. We used these two independent instances of colonization and results from a rearing experiment to quantify the degree to which genomic changes following alfalfa colonization were predictable. We found a modest overlap in SNPs showing the greatest allele frequency differences between alfalfa and native-feeding *L. melissa* in the western and eastern populations (~1.5-2.8 times more than expected by chance, depending on the quantile considered), and a significant but weaker and more idiosyncratic excess in overlap of SNPs associated with host-specific larval performance in an experiment and those with the greatest allele frequency differences between host-associated populations in nature (~0.53-2.5 times more than expected by chance, depending on the quantile considered and performance measure). Although we were able to predict (to a modest extent) the SNPs with the greatest genomic change in nature (in terms of parallelism in nature and from the experimental results), we generally had little to no ability to predict the direction of change (i.e., even if the same SNP showed exceptional change in *melissa-east* and melissa-west, the direction of change was not necessarily the same). SNP × performance associations were, however, more predictive of patterns of genetic differentiation in nature between the specific populations used in the rearing and mapping experiments. We discuss and interpret these results in more detail below.

### Predictability of genomic changes associated with a host shift

We identified a significant excess of shared SNPs between *melissa-east* and *melissa*-west populations, specifically ~ 1.5-2.8 times more than expected by chance. This means that knowing which SNPs exhibited the greatest genomic change in one geographic group (i.e., in one colonization event) improves our ability to predict those with the greatest genomic change in the other group about two-fold. In some ways, such predictability is not surprising as the western and eastern populations likely had access to much of the same standing genetic variation (Conte *et al.,* 2012). But there are also reasons to think that parallelism at the genetic level (and thus predictability of genomic change) might be more limited. For example, alfalfa is not a homogeneous resource, and our previous work has documented variation in caterpillar performance based on the source of alfalfa (Harrison *et al.,* 2016), which suggests that the way in which a population adapts to alfalfa might depend on the specific host plant population. Interestingly, a nearly identical excess of parallel genomic change/genetic differentiation was detected in comparisons of host-associated stick insects *(Timema cristinae*) where different cryptic color patterns are favored on different hosts (Soria-Carrasco *et al.,* 2014). For simpler morphological traits, such as armor plating in sticklebacks, the degree of parallelism is considerably higher (91% of the high genetic differentiation regions shared between marine and freshwater populations across population pairs are also shared between additional populations) (Jones *et al.,* 2012). Another study in sticklebacks shows high parallel allele frequency changes between lake and stream ecotype populations in genomic regions associated with incipient ecological speciation ( 51% of the genomic islands of differentiation show parallel changes) (Marques *et al.,* 2016).

SNP × performance associations in lab experiments predicted genomic change in nature, but in a more limited and more idiosyncratic way; x-fold enrichments for survival range from 0.63-1.95 and x-fold enrichments for weight range from 0.53-2.5, with only ~25% of combinations showing significant excess. Predictability was notably higher in terms of predicting the most differentiated SNPs between the populations used in the rearing experiment, that is, GLA (host = *M. sativa*) and SLA (host = *A. canadensis*); the x-fold enrichment ranged from 1.42 to 4.53 (across comparisons), with all but one case significant. Given the simplified lab rearing environment (e.g., no predators, controlled growth conditions, only some fitness components considered, etc.), it is intriguing that the experiment provided even these level of predictive power about genomic change in nature. With that said, these results are consistent with two other recent studies that predicted genomic change in nature from short-term experiments. In *Timema cristinae* stick insects, Soria-Carrasco *et al.* (2014) found a modest but significant overlap between genetic regions associated with survival in a field experiment and those most differentiated between hosts in nature (obs. = 32 shared loci; expected = 23; x-fold enrichment = 1.4). In a similar study with *Rhagoletis pomonella* fruit flies, genomic change during a lab selection experiment was even more predictive of patterns of differentiation in nature (obs. = 154 shared loci; expected = 53.6; x-fold = 2.87) (Egan *et al.,* 2015). Substantial genomic change and increased predictability in *Rhagoletis* might be due, at least in part, to the high levels of LD in that system.

Beyond host-use in herbivorous insects, a limited number of studies have tried to predict genome-wide patterns of genetic differentiation in nature from lab or field experiments, and mostly these have involved predicting genetic differentiation from QTL studies. The outlier loci underlying highest genetic differentiation between *Drosophila yakuba* mainland (Cameroon and Kenya) and Mayotte populations, show concordance with *Drosophila sechellia* noni-tolerance (performance) QTLs (four of nine tolerance QTL, expected = 0.125, P = 0.013) but there is no overlap for *D. sechellia* preference QTLs (expected = 0.35, P = 0.37), suggesting that noni-performance is more predictable than noni preference (Yassin *et al.,* 2016; Vertacnik & Linnen, 2017). Similarly, QTL for ecologically relevant traits co-localized with possible genetic regions affected by selection (as identified in genome scans) more than expected by chance in comparisons of lake whitefish ecotypes (Rogers & Bernatchez, 2007). The predictability of evolution from lab selection and mapping experiments will depend on whether the same genetic variants affect key traits in similar ways in the lab and nature. This has been investigated in several taxa. For example, in *Arabidopsis thaliana,* only one QTL associated with flowering time in the greenhouse also affected flowering time in a field experiment meant to better approximate nature (Weinig *et al.,* 2002; Brachi *et al.,* 2010).

Despite results suggesting the SNPs with the greatest genomic change during host adaptation could be predicted, both via parallel change in nature and from a short-term rearing/mapping experiment, we found much less evidence that the direction of genomic change was predictable (there were a few, limited exceptions). Even if the same alleles are repeatedly favored on alfalfa (including in the lab), the direction of change at genetic markers could vary if patterns of LD between sequenced SNPs and causal variants differ. For example, if the favored allele at a causal locus is positively associated with one SNP marker allele in one population and negatively associated with the same SNP marker allele in another population, selection on the causal locus could drive substantial evolutionary change at the SNP locus in both populations, but in opposite directions. Such shifts in patterns of LD could occur when new populations are founded (possibly by one or a few gravid females), and thus, this phenomenon could explain our results. Consistent with this possibility, we found slightly more cases (18.8% vs. 14.6% of tests with P ≤ 0.05) of excess sign coincidence when comparing the experimental results to genetic differentiation between GLA and SLA than when comparing them to overall patterns of SNP × host-use association. In a related sense, if recombination rates vary across the genome, regions of exceptional genomic change might be predictable if they are simply the regions with the lowest recombination rate and thus the lowest local effective population size. Of course, the rate of evolutionary change by drift or selection is proportional to the allele frequencies (specifically to *p*(1 ― *p*)), which could make the regions of greatest change (but not their direction) predictable, but this is something we have controlled for by working with the residuals after regressing change (or effect sizes from the rearing experiment) on allele frequencies.

In contrast to our results, both the magnitude and direction of genomic change between host races of *Rhagoletis pomonella* fruit flies was predictable from a controlled experiment (Egan *et al.,* 2015). Two factors likely contribute to this difference. First, the same populations were used in the experiment and for the natural comparisons, whereas our experiment focused on a small subset of the natural populations we analyzed (but see the discussion of the comparison with genetic differentiation between GLA and SLA above). Second, inversions and large blocks of LD are common in *Rhagoletis* and could increase the consistency of evolutionary patterns and associations between SNP markers and causal variants. Although not concerned with host adaptation, another study which has tested for direction of allele frequency changes underlying rapid adaptation focuses on adaptation to fragmented landscapes in Glanville Fritillary butterflies, *Melitaea cinxia* (Fountain *et al.,* 2016). This study reports predictable allele frequency changes in most divergent outlier loci between newly colonized versus old local populations, and these allele frequency shifts are in the same direction indicating that selection can drive particular candidate genetic regions in the direction of adaptation to fragmented landscapes (Extinct populations: linear model, *r*^2^ = 0.36, *P* = 0.02; Introduced populations: linear model, *r*^2^ = 0.14, *P* = 0.13).

Finally, we found higher predictability of genomic change in terms of the greater overlap of top host-associated SNPs between two host shifts onto alfalfa than overlap between performance-associated SNPs in an experiment and host-associated SNPs in nature (predictability in terms of overlap was even higher for patterns of genetic differentiation between the population pair used in the experiment). Perhaps this is not surprising, as we might expect greater similarity in conditions across the natural populations than between the lab experiment and natural populations. Indeed, perhaps it is more surprising that the difference in predictability (1.5-2.7 x-fold enrichment via parallelism in nature vs. 0.53-2.5 x-fold enrichment via predictions from the lab to nature) wasn’t greater. This means that genetic and phenotypic determinants of caterpillar performance in the lab have some bearing on fitness, and thus evolutionary change during host shifts in nature. Posing the exciting possibility that, by integrating outcomes from multiple experiments probing different populations and different components of a host shift (e.g., larval performance, adult preference, etc.), it might be possible to build a more mechanistic model to predict genomic change than would ever be possible by only examining patterns of parallel change in nature.

### Interpretation of demographic patterns

We found little evidence for contemporary gene flow among *L. melissa* populations, even those separated by only a few kilometers. This suggests that gene flow from populations feeding on native hosts to populations feeding on alfalfa is not a major factor constraining host adaptation. Moreover, coupled with our previous results indicating a lack of genetic trade-offs for larval performance across hosts (Gompert *et al.,* 2014a), this suggests that host plant specialization in *L. melissa* could occur via the loss of adaptation to an ancestral host by genetic drift. Similarly, a recent study in moths *(Thyrinteina leucoceraea*) found that the loss of adaptation to a native host was due to mutation accumulation rather than trade-offs (Grosman *et al.,* 2015). Thus, whereas resource-based genetic trade-offs do drive host specialization in some systems (Gompert & Messina, 2016), this and other recent work indicates that other processes that need not include selection can lead to host-plant specialization as well.

Our results strongly suggest that *L. melissa* colonized alfalfa multiple times since the introduction of this plant to North America; at least twice and probably ~four times. Shifts from *M. sativa* to native hosts appear to be even more common, which is consistent with our own observations that populations associated with *M. sativa* are more ephemeral (i.e., less likely to persist over multiple decades) than those feeding on native hosts. Still, considerable uncertainty exists in our estimates of the exact number and nature of these host shifts. Along these lines, more than one colonization event likely occurred within our eastern and western *L. melissa* groups. Thus, while treating these groups as our level of replication is conservative, it also means that the metrics of parallelism discussed above probably represent averages over these putative additional host shifts, and that the true history of colonization is more complex than captured in these analyses.

### Genomic context of the host-associated SNPs

An excess of the SNPs most associated with host use in nature were on the Z chromosome in *L. melissa.* This is consistent with findings from other studies suggesting a disproportionate role for sex chromosomes in adaptation and speciation, such as beak morphology in Darwin’s finches (Abzhanov *et al.,* 2004, 2006), coat color in mice *(Chaetodipus intermedins*) (Nachman *et al.,* 2003; Hoekstra *et al.,* 2006), reduction in armor plating in threespine sticklebacks *(Gasterosteus aculeatus*) (Colosimo *et al.,* 2005), and wing pattern variation in *Heliconious* butterflies (Papa *et al.,* 2008). In butterflies, other studies of host plant specialization have found evidence of a disproportionate role of Z-linked genes in host specificity and oviposition preference (Matsubayashi *et al.,* 2010). For example, oviposition preference in the comma butterfly *(Polygonia c-album*) is Z-linked (Janz, 1998; Nygren *et al.,* 2006). Similarly, oviposition differences in some swallowtail butterflies *(Papilio zelicaon* and *Papilio oregonius*) are known to be sex-linked (Thompson, 1988), and in general, species differences in butterflies often map to the Z chromosome (Sperling, 1994; Prowell, 1998). With that said, it is important to note that the Z chromosome likely has a lower effective population size than the autosomes (this depends some on patterns of mating), and thus the signal of an excess of host-associated SNPs on the Z chromosome could partially reflect genetic drift.

We also found an excess of top host-associated SNPs on the coding regions of genes (1.45 times more than expected by chance). This does not necessarily imply a greater role for structural (vs. regulatory) changes in host adaptation, as these SNPs could also be in LD with nearby regulatory elements. But it does bolster the evidence that these SNPs are tagging (via LD) some causal variants (i.e., that their status as top host-associated SNPs does not solely reflect a greater role for genetic drift). Similar results have been seen in several other genome scans for selection or adaptation (Counterman *et al.,* 2010; Jones *et al.,* 2012; Soria-Carrasco *et al.,* 2014). Finally, we found that the genes nearest to the top host-associated SNPs have annotations suggesting a diversity of molecular functions and biological/cellular processes (Table S4). None of these stands out in a clear way, but this does suggest that host adaptation is likely a multifaceted process with selection shaping many different genes and molecular or developmental pathways.

## Acknowledgements

This research was supported by the National Science Foundation (DEB-1638768 to ZG, DEB-1050355 to CCN and DEB-1050726 to MLF) and by the Utah Agricultural Experiment Station (approved as journal paper number XXXX). The support and resources from the Center for High Performance Computing at the University of Utah are gratefully acknowledged.

## Data Accessibility

DNA sequence data will be archived in the NCBI SRA (Submission ID: SUB3611688, BioProject ID: PRJNA432816). The genome sequence, annotations, sequence alignments, variant data, baypass infiles, and other analyses results and computer scripts will be archived in DRYAD (doi:10.5061/dryad.87ht5v0).

## Author Contributions

ZG, MLF and CCN conceived of the project. LKL, ZG, and CCN generated the data. SC and ZG analyzed the data. SC and ZG wrote the manuscript. All authors helped revise the

## Supplemental tables and figures

**Table S1:** Table shows summary of randomization tests for top 0.01% host-associated SNPs and top 0.01% parallel host-associated SNPs for presence on Z-chromosome (No. observed = number of SNPs observed on the sex chromosome; x-fold = number of observed is how much more than chance; number of SNPs observed on Z-chromosome and tests for randomizations; P = randomization-based P-values for the null hypothesis that the proportion of top SNPs observed on the Z-chromosome is not greater than the genomic proportion).

| Set | No. observed | P | *x-fold* |
| --- | --- | --- | --- |
| *All populations* |  |  |  |
| top 0.01% host-associated | 195 | <**0.01** | 2.26 |
| top 0.01% parallelism | 6 | 0.03 | 2.48 |
| *melissa-east* |  |  |  |
| top 0.01% host-associated | 193 | <**0.01** | 2.23 |
| top 0.01% parallelism | 6 | 0.03 | 2.48 |
| *melissa-west* |  |  |  |
| top 0.01% host-associated | 134 | <**0.01** | 1.55 |
| top 0.01% parallelism | 6 | 0.04 | 2.48 |

**Table S2:** Table shows summary of randomization tests for presence of top (0.01%) host-use associated SNPs on gene region of the genome (Top SNP% = Quantiles cut off for analysis; x-fold = Number of SNPs observed is how much more than chance; P = randomization-based P-values for the null hypothesis that the proportion of top SNPs observed is not greater than the genomic proportion; Mean = mean for the null hypothesis). P ≤ 0.05 are in bold.

| Top SNP % | No. observed | $P$ | x-fold enrichment |
| --- | --- | --- | --- |
| *All populations* |  |  |  |
| top 0.001% | 0.13 | **0.01** | 1.30 |
| top 0.01% | 0.42 | 0.82 | 0.96 |
| top 0.1% | 0.34 | 1.00 | 0.51 |
| *melissa-east* |  |  |  |
| top 0.001% | 0.15 | < **0.01** | 1.30 |
| top 0.01% | 0.42 | 0.86 | 0.96 |
| top 0.1% | 0.31 | 0.26 | 1.01 |
| *melissa-west* |  |  |  |
| top 0.001% | 0.14 | < **0.01** | 1.30 |
| top 0.01% | 0.47 | 0.91 | 0.96 |
| top 0.1% | 0.32 | 1.00 | 0.51 |
| *top 0.01% parallelism* |  |  |  |
| top 0.001% | 0 | 0.99 | 0.5 |
| top 0.01% | 0 | 1.00 | 0 |

**Table S3:** Table shows summary of randomization tests for presence of top (0.01%) host-use associated SNPs on coding region of the genome (To SNP% = quantiles cut off for analysis; x-fold = Number of SNPs observed is how much more than chance; P = randomization-based P-values for the null hypothesis that the proportion of top SNPs observed is not greater than the genomic proportion; Mean = mean for the null hypothesis). P significant at 0.05 are in bold.

| Set | *No. observed* | P | x-fold |
| --- | --- | --- | --- |
| *All populations* |  |  |  |
| top 0.001% | 1.00 | 0.12 | 0.54 |
| top 0.01% | < **0.01** | 0.43 | 1.46 |
| top 0.1% | < **0.01** | 0.31 | 1.07 |
| *melissa-east* |  |  |  |
| top 0.001% | 1.00 | 0.14 | 0.54 |
| top 0.01% | < **0.01** | 0.42 | 1.46 |
| top 0.1% | < **0.01** | 0.30 | 1.07 |
| *melissa-west* |  |  |  |
| top 0.001% | 1.00 | 0.14 | 0.54 |
| top 0.01% | < **0.01** | 0.42 | 1.46 |
| top 0.1% | < **0.01** | 0.34 | 1.07 |
| *top 0.01% parallelism* |  |  |  |
| top 0.001% | 0.99 | < **0.01** | 0.51 |
| top 0.01% | 1.00 | < **0.01** | 0.00 |

**Table S4:** Table shows summary of randomization tests for determining molecular functions of top (0.01%) host-use associated SNPs (GO ID = Gene ontology reference ID; GO name = Name of the function associated with GO ID, No. = number of top 0.01% SNPs enriched for the GO function, P = randomization-based *P*-values for the null hypothesis that the proportion of top SNPs observed is not greater than the genomic proportion, x-fold = number of SNPs observed is how much more than chance. P ≤ 0.05 are in bold.

| Molecular functions |  |  |  |  |
| --- | --- | --- | --- | --- |
| GO ID | GO name | No. | $P$ | x-fold |
| GO:0004519 | endonuclease activity | 8 | < **0.01** | 4.92 |
| GO:0004177 | aminopeptidase activity | 7 | **0.01** | 2.41 |
| GO:0008234 | Cysteine-type peptidase activity | 7 | **0.04** | 2.25 |
| GO:0003684 | damaged DNA binding | 7 | < **0.01** | 6.23 |
| GO:0005452 | inorganic anion exchanger activity | 6 | **0.04** | 2.16 |
| GO:0004047 | aminomethyltransferase activity | 6 | < **0.01** | 2.92 |
| GO:0005272 | sodium channel activity | 5 | **0.01** | 2.68 |
| GO:0008889 | glycerophosphodiester phosphodiesterase activity | 5 | **0.05** | 2.12 |
| GO:0005544 | Calcium-dependent phospholipid binding | 4 | **0.04** | 2.33 |
| GO:0004768 | Stearoyl-CoA 9-desaturase activity | 4 | 0.06 | 2.25 |
| GO:0004044 | amidophosphoribosyltransferase activity | 4 | **0.01** | 2.79 |
| GO:0016709 | oxidoreductase activity* | 4 | **0.01** | 3 |
| GO:0005247 | Voltage-gated chloride channel activity | 3 | 0.08 | 2.32 |
| GO:0033897 | ribonuclease T2 activity | 3 | **0.02** | 3.61 |
| GO:0004452 | Isopentenyl-diphosphate delta-isomerase activity | 3 | 0.12 | 2.56 |
| GO:0030429 | kynureninase activity | 3 | 0.13 | 2.54 |
| GO:0016844 | strictosidine synthase activity | 3 | < **0.01** | 4.23 |
| GO:0016538 | Cyclin-dependent protein serine/threonine kinase regulator activity | 3 | 0.14 | 2.17 |
| GO:0003867 | 4-aminobutyrate transaminase activity | 3 | **0.02** | 3.32 |
| GO:0004594 | pantothenate kinase activity | 2 | **0.04** | 3.34 |
| GO:0017172 | cysteine dioxygenase activity | 2 | **0.01** | 4.91 |
| GO:0005201 | extracellular matrix structural constituent | 2 | **0.04** | 2.84 |
| GO:0008565 | protein transporter activity | 2 | 0.12 | 2.48 |
| GO:0004066 | asparagine synthase (glutamine-hydrolyzing) activity | 2 | **0.02** | 3.33 |
| GO:0003950 | NAD+ ADP-ribosyltransferase activity | 2 | **0.02** | 4.14 |

**Table S5:** Table shows summary of randomization tests for determining biological functions of top (0.01%) host-use associated SNPs (GO ID = Gene ontology reference ID; GO name = Name of the function associated with GO ID, No. = Number of top 0.01% SNPs enriched for the GO function, P = randomization-based *P*-values for the null hypothesis that the proportion of top SNPs observed is not greater than the genomic proportion, x-fold = Number of SNPs observed is how much more than chance. P ≤ 0.05 are in bold.

| Biological functions |  |  |  |  |
| --- | --- | --- | --- | --- |
| GO ID | GO name | No. | $P$ | x-fold |
| GO:0006741 | NADP biosynthetic process | 9 | **0.01** | 2.13 |
| GO:0019674 | NAD metabolic process | 9 | **0.01** | 2.15 |
| GO:0015991 | ATP hydrolysis coupled proton transport | 9 | **0.02** | 2.14 |
| GO:0006546 | glycine catabolic process | 6 | **0.01** | 2.68 |
| GO:0006820 | anion transport | 6 | **0.04** | 2.12 |
| GO:0006071 | glycerol metabolic process | 5 | **0.04** | 2.31 |
| GO:0019752 | carboxylic acid metabolic process | 5 | **0.02** | 2.54 |
| GO:0015937 | coenzymeA biosynthetic process | 5 | 0.06 | 2.77 |
| GO:0006542 | glutamine biosynthetic process | 5 | **0.05** | 2.32 |
| GO:0042318 | penicillin biosynthetic process | 4 | 0.12 | 2.23 |
| GO:0006506 | GPI anchor biosynthetic process | 4 | 0.14 | 2.21 |
| GO:0009116 | nucleoside metabolic process | 4 | **0.03** | 2.81 |
| GO:0009113 | purine nucleobase biosynthetic process | 4 | **0.02** | 2.83 |
| GO:0006633 | fatty acid biosynthetic process | 4 | 0.09 | 2.32 |
| GO:0006744 | ubiquinone biosynthetic process | 4 | **0.01** | 2.91 |
| GO:0006821 | chloride transport | 3 | 0.07 | 2.23 |
| GO:0009448 | gamma-aminobutyric acid metabolic process | 3 | **0.02** | 3.23 |
| GO:0006397 | mRNA processing | 3 | **0.03** | 2.91 |
| GO:0006569 | tryptophan catabolic process | 3 | 0.12 | 2.44 |
| GO:0030071 | regulation of mitotic metaphase/anaphase transition | 2 | **0.04** | 3.21 |
| GO:0006904 | vesicle docking involved in exocytosis | 2 | **0.02** | 3.92 |
| GO:0046439 | L-cysteine metabolic process | 2 | **0.01** | 4.95 |
| GO:0042254 | ribosome biogenesis | 2 | 0.13 | 2.1 |
| GO:0000398 | mRNA splicing via spliceosome | 2 | **0.03** | 3.25 |
| GO:0006529 | asparagine biosynthetic process | 2 | **0.04** | 3.22 |

**Table S6:** Table shows summary of randomization tests for determining cellular functions of top (0.01%) host-use associated SNPs (GO ID = Gene ontology reference ID; GO name = Name of the function associated with GO ID, No. = Number of top 0.01% SNPs enriched for the GO function, P = randomization-based *P*-values for the null hypothesis that the proportion of top SNPs observed is not greater than the genomic proportion, x-fold = Number of SNPs observed is how much more than chance. P ≤ 0.05 are in bold.

| Cellular functions |  |  |  |  |
| --- | --- | --- | --- | --- |
| GO ID | GO name | No. | $P$ | x-fold |
| GO:0005643 | nuclear pore | 10 | < **0.01** | 2.33 |
| GO:0000139 | Golgi membrane | 6 | **0.01** | 2.72 |
| GO:0005795 | Golgi stack | 4 | < **0.001** | 9.61 |
| GO:0033180 | proton-transporting V-type ATPase V1 domain | 2 | **0.01** | 4.83 |
| GO:0005581 | collagen trimer | 2 | **0.05** | 2.75 |
| GO:0030126 | COPI vesicle coat | 2 | **0.02** | 3.24 |
| GO:0005960 | glycine cleavage complex | 2 | **0.01** | 4.81 |

**Table S7:** Results from randomization tests for overlap of the top (0.01%) host-use associated SNPs in nature and the top (0.01%) survival-associated SNPs in rearing experiment (based on *ran1*). Population-plant = population and plant treatment in the laboratory experiment; No. observed = number of SNPs associated with both host use in wild and performance in the lab; x-fold = enrichment relative to null expectations; P = randomization-based P-values for the null hypothesis (P ≤ 0.05 are in bold). Results are shown based on raw Bayes factors and model-averaged effect sizes, and based on residuals controlling these metrics for allele frequencies.

| Population-plant | Raw values |  |  | Residual values |  |  |
| --- | --- | --- | --- | --- | --- | --- |
| | No. observed | $P$ | x-fold | No. observed | $P$ | x-fold |
| *All populations* |  |  |  |  |  |  |
| GLA-Ms | 4 | 0.96 | 0.49 | 14 | **0.03** | 1.74 |
| GLA-Ac | 6 | 0.84 | 0.72 | 11 | 0.21 | 1.34 |
| SLA-Ms | 2 | 0.99 | 0.25 | 9 | 0.39 | 1.14 |
| SLA-Ac | 0 | 1.00 | 0.00 | 3 | 0.39 | 0.98 |
| *melissa-east* |  |  |  |  |  |  |
| GLA-Ms | 6 | 0.81 | 0.74 | 13 | 0.06 | 1.62 |
| GLA-Ac | 7 | 0.71 | 0.85 | 16 | **0.01** | 1.95 |
| SLA-Ms | 2 | 0.99 | 0.25 | 11 | 0.17 | 1.39 |
| SLA-Ac | 4 | 0.95 | 0.52 | 9 | 0.36 | 1.17 |
| *melissa-west* |  |  |  |  |  |  |
| GLA-Ms | 9 | 0.41 | 1.12 | 13 | 0.07 | 1.61 |
| GLA-Ac | 13 | 0.07 | 1.58 | 14 | **0.04** | 1.71 |
| SLA-Ms | 4 | 0.95 | 0.50 | 5 | 0.89 | 0.63 |
| SLA-Ac | 4 | 0.94 | 0.52 | 9 | 0.35 | 1.18 |

**Table S8:** Results from randomization tests for overlap of the top (0.01%) host-use associated SNPs in nature and the top (0.01%) weight-associated SNPs in rearing experiment (based on *ran1*). Population-plant = population and plant treatment in the laboratory experiment; No. observed = number of SNPs associated with both host use in wild and performance in the lab; x-fold = enrichment relative to null expectations; P = randomization-based P-values for the null hypothesis (P ≤ 0.05 are in bold). Results are shown based on raw Bayes factors and model-averaged effect sizes, and based on residuals controlling these metrics for allele frequencies.

| Population-plant | Raw values |  |  | Residual values |  |  |
| --- | --- | --- | --- | --- | --- | --- |
| | No. observed | $P$ | x-fold | No. observed | $P$ | x-fold |
| *All populations* |  |  |  |  |  |  |
| GLA-Ms | 5 | 0.89 | 0.64 | 11 | 0.16 | 1.42 |
| GLA-Ac | 21 | < **0.01** | 2.67 | 11 | 0.17 | 1.39 |
| SLA-Ms | 22 | < **0.01** | 2.87 | 10 | 0.23 | 1.31 |
| SLA-Ac | 2 | 0.99 | 0.26 | 6 | 0.78 | 0.78 |
| *melissa-east* |  |  |  |  |  |  |
| GLA-Ms | 7 | 0.66 | 0.90 | 8 | 0.52 | 1.03 |
| GLA-Ac | 5 | 0.89 | 0.64 | 10 | 0.25 | 1.28 |
| SLA-Ms | 5 | 0.95 | 0.52 | 4 | 0.95 | 0.53 |
| SLA-Ac | 8 | 0.49 | 1.05 | 17 | < **0.01** | 2.23 |
| *melissa-west* |  |  |  |  |  |  |
| GLA-Ms | 16 | **0.01** | 2.08 | 16 | **0.01** | 2.06 |
| GLA-Ac | 12 | 0.09 | 1.53 | 13 | **0.05** | 1.66 |
| SLA-Ms | 10 | 0.23 | 1.32 | 19 | < **0.01** | 2.50 |
| SLA-Ac | 10 | 0.24 | 1.31 | 12 | 0.09 | 1.56 |

**Table S9:** Table shows summary of randomization tests for concordance in effect signs of overlapping host-associated SNPs and survival-associated SNPs in the rearing experiment for the top 0.01% empirical quantile (P ≤ are in bold). Results are shown for randomization tests *ran2A* and *ran2B.*

| Population-plant | No. observed | *ran2A* |  | *ran2B* |  |
| --- | --- | --- | --- | --- | --- |
|  |  | *P* | x-fold | *P* | x-fold |
| *All populations* |  |  |  |  |  |
| GLA-Ms | 8 | 0.16 | 1.07 | 0.23 | 1.11 |
| GLA-Ac | 4 | 0.39 | 0.87 | 0.69 | 0.75 |
| SLA-Ms | 5 | 1.00 | 1.00 | 0.27 | 1.08 |
| SLA-Ac | 1 | 0.36 | 0.58 | 0.48 | 0.67 |
| *melissa-east* |  |  |  |  |  |
| GLA-Ms | 9 | **0.05** | 1.21 | **0.04** | 1.38 |
| GLA-Ac | 5 | 0.81 | 0.64 | 0.88 | 0.64 |
| SLA-Ms | 9 | **< 0.01** | 1.39 | **0.01** | 1.59 |
| SLA-Ac | 4 | 1.00 | 1.00 | 0.49 | 0.89 |
| *melissa-west* |  |  |  |  |  |
| GLA-Ms | 7 | 0.54 | 0.85 | 0.29 | 1.07 |
| GLA-Ac | 9 | 0.28 | 1.01 | 0.36 | 1.03 |
| SLA-Ms | 4 | **< 0.01** | 1.42 | **0.03** | 1.53 |
| SLA-Ac | 5 | 0.12 | 1.15 | 0.26 | 1.11 |

**Table S10:** Table shows summary of randomization tests for concordance in effect signs of overlapping host-associated SNPs and weight-associated SNPs in the rearing experiment for the top 0.01% empirical quantile (significant P-values at 0.05 are in bold). Results are shown for randomization tests *ran2A* and *ran2B.*

| Population-plant | No. observed | *ran2A* |  | *ran2B* |  |
| --- | --- | --- | --- | --- | --- |
|  |  | *P* | x-fold | *P* | x-fold |
| *All populations* |  |  |  |  |  |
| GLA-Ms | 7 | 0.13 | 1.07 | 0.13 | 1.25 |
| GLA-Ac | 5 | 0.45 | 0.85 | 0.46 | 0.92 |
| SLA-Ms | 2 | 0.71 | 0.58 | 0.93 | 0.41 |
| SLA-Ac | 1 | 0.66 | 0.43 | 0.87 | 0.34 |
| *melissa-east* |  |  |  |  |  |
| GLA-Ms | 4 | 0.17 | 1.07 | 0.36 | 1.02 |
| GLA-Ac | 2 | 0.94 | 0.39 | 0.93 | 0.41 |
| SLA-Ms | 1 | 0.48 | 0.51 | 0.69 | 0.49 |
| SLA-Ac | 2 | 0.99 | 0.25 | 0.99 | 0.24 |
| *melissa-west* |  |  |  |  |  |
| GLA-Ms | 7 | 0.61 | 0.82 | 0.59 | 0.87 |
| GLA-Ac | 5 | < **0.01** | 1.21 | 0.68 | 0.78 |
| SLA-Ms | 7 | 0.59 | 0.85 | 0.82 | 0.74 |
| SLA-Ac | 6 | 0.35 | 0.93 | 0.39 | 1.01 |

**Table S11:** Table shows summary of randomization tests for overlapping high F_ST_ SNPs and performance-associated SNPs in the rearing experiment for the top 0.01% empirical quantile (P ≤ 0.05 are in bold). Results are shown for randomization tests *ran1*

| Population-plant | No. observed | $P$ | x-fold |
| --- | --- | --- | --- |
| *GLA-SLA Pairwise $F_{ST}$* |  |  |  |
| GLA-Ms-weight | 20 | **<0.01** | 2.58 |
| GLA-Ac-weight | 14 | **0.02** | 1.81 |
| GLA-Ms-survival | 14 | **0.02** | 1.81 |
| GLA-Ac-survival | 29 | **<0.01** | 3.75 |
| SLA-Ms-weight | 21 | **< 0.01** | 2.71 |
| SLA-Ac-weight | 35 | **< 0.01** | 4.53 |
| SLA-Ms-survival | 13 | **0.04** | 1.68 |
| SLA-Ac-survival | 11 | 0.16 | 1.42 |
| *GLA-ABC Pairwise $F_{ST}$* |  |  |  |
| GLA-Ms-weight | 12 | 0.09 | 1.55 |
| GLA-Ac-weight | 13 | **0.05** | 1.68 |
| GLA-Ms-survival | 15 | **0.01** | 1.93 |
| GLA-Ac-survival | 13 | **0.04** | 1.68 |
| *SLA-VCP Pairwise $F_{ST}$* |  |  |  |
| SLA-Ms-weight | 13 | **0.05** | 1.67 |
| SLA-Ac-weight | 16 | **0.01** | 2.06 |
| SLA-Ms-survival | 6 | 0.79 | 0.77 |
| SLA-Ac-survival | 7 | 0.65 | 0.91 |

**Table S12:**
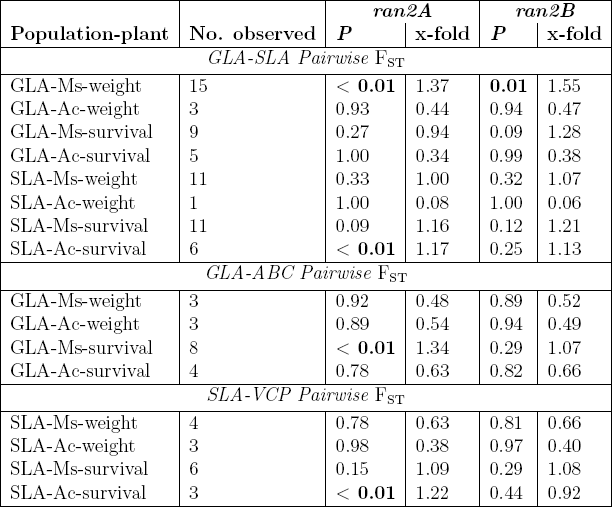
Table shows summary of randomization tests for concordance in effect signs of overlapping pairwise high F_ST_ SNPs and performance-associated SNPs in the rearing experiment for the top 0.01% empirical quantile (P ≤ 0.05 are in bold). Results are shown for randomization tests *ran2A* and *ran2B.*

**Figure S1:**
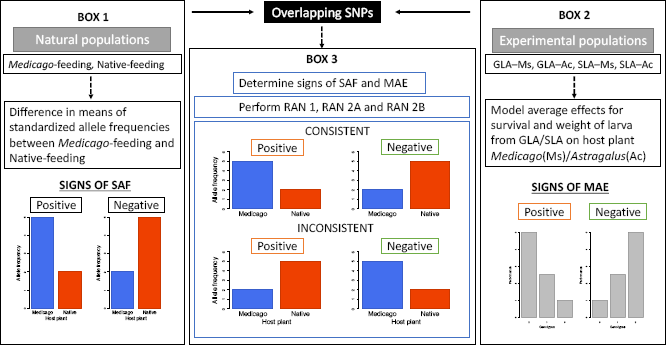
Diagram shows a schematic representation of the analyses conducted to test for concordance between direction of allele frequency differences between alfalfa-feeding and native-feeding populations and signs for model average effects for performance-associated SNPs in rearing experiment. Each box represents an analysis conducted in the study SAF = standardized allele frequencies for host-associated SNPs in natural populations, MAE = model average effects for performance-associated SNPs.

**Figure S2:**
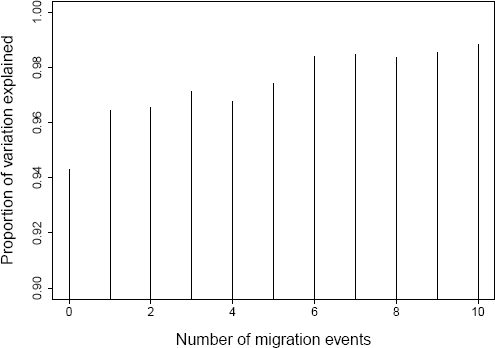
Plot shows proportion of variation explained by the `TREEMIX` population graph with different numbers of migration edges.

**Figure S3:**
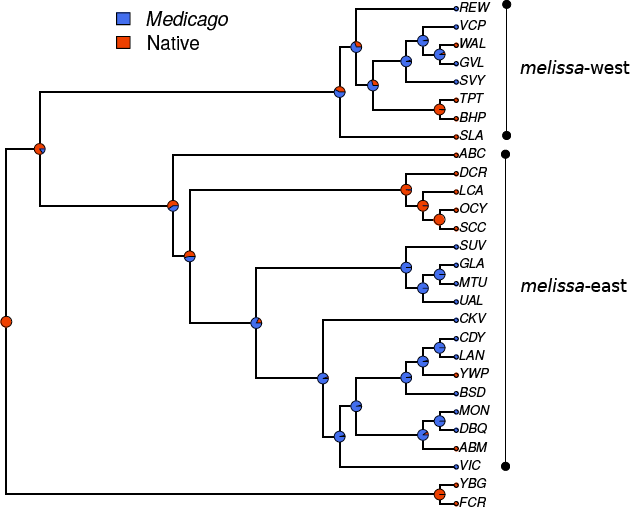
Tree shows ancestral state recontruction of the mutations that lead to colonization shifts from native host to novel host *Medicago sativa.* Terminal nodes are labeled by abbreviations for geographical locations from where samples were collected and circles beside the terminal locations are colored according to host-plant association. Inferred ancestral states are denoted by pie-charts that indicate the posterior probability of being associated with native host (orangered) versus being associated with *Medicago* (blue).

**Figure S4:**
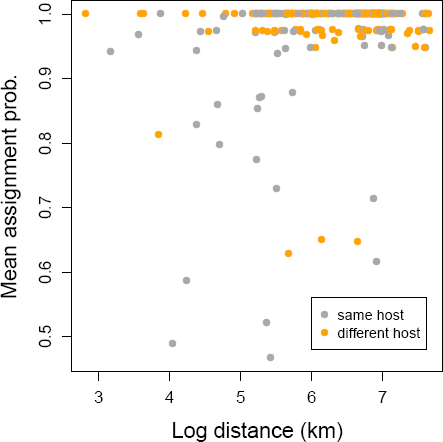
Plot shows the mean assignment probability to the correct population (i.e., the one that an individual was sampled from) across all 300 pairs as a function of log geographic distance and whether the pair of populations feed on the same or different host plants. Note that average assignments to the collected populations were very similar for same (0.964, sd = 0.0524) and different (0.984, sd = 0.953) host comparisons.

**Figure S5:**
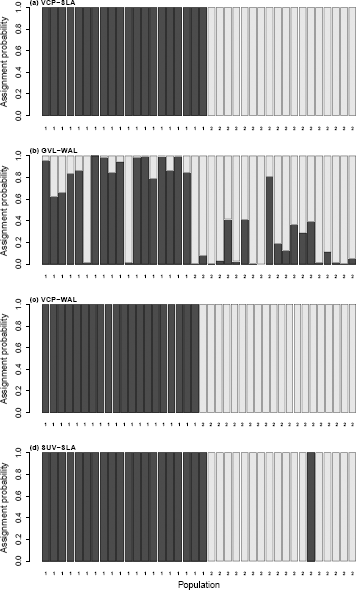
Barplots show individual assignment probabilities for four of the nearest population pairs that fed on different host plants. In panels (A) and (C) all individuals were confidently assign to the population they were collected from. (B) shows a case where that there is much more uncertainty in general (i.e., genetic differentiation between these populations is low), but two likely migrants. (D) shows a single individual that is most likely a migrant from SUV (or a similar population) to SLA.

**Figure S6:**
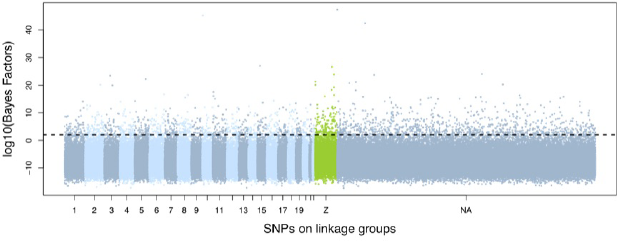
Manhattan plot for all populations (N=25) shows SNPs (N=206,028) as points mapped along linkage groups (1-Z). Z indicates the sex-chromosome. NA indicates SNPs which have not been assigned to a linkage group. Straight line separates the top 0.01% SNPs with high Bayes factor values.

**Figure S7:**
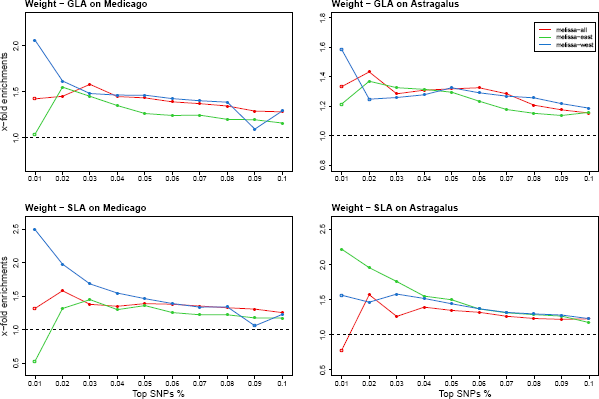
Line plots show x-fold enrichments across quantiles for overlapping SNPs between weight-associated SNPs in the rearing experiment and host-associated SNPs in nature. Open circles indicate P > 0.05 and filled circles indicate P < 0.05.

**Figure S8:**
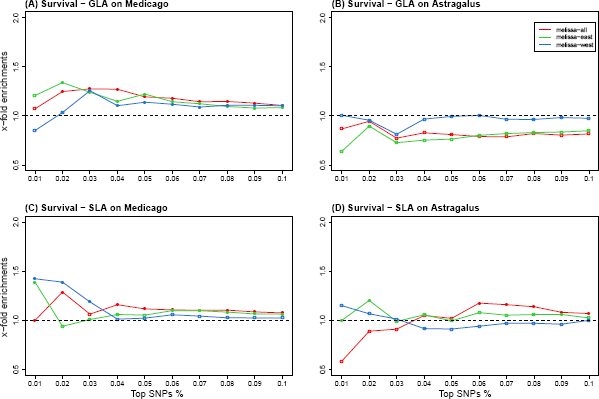
Line plot shows x-fold enrichments across quantiles for overlapping SNPs between survival-associated SNPs in rearing experiment and host-associated SNPs in nature for *ran2A*. Open circles indicate P > 0.05 and filled circles indicate P < 0.05.

**Figure S9:**
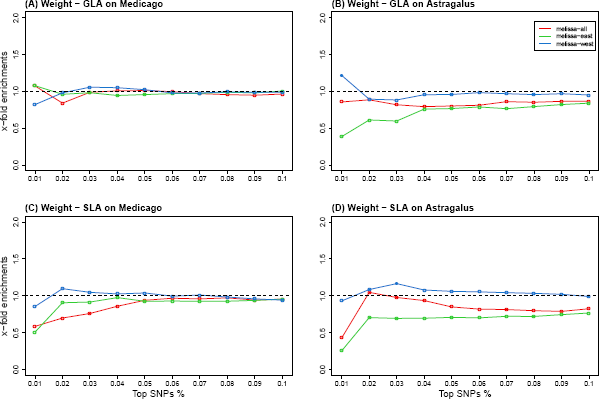
Line plots show x-fold enrichments across quantiles for overlapping SNPs between weight-associated SNPs in the rearing experiment and host-associated SNPs in nature for *ran2A*. Open circles indicate P > 0.05 and filled circles indicate P < 0.05.

**Figure S10:**
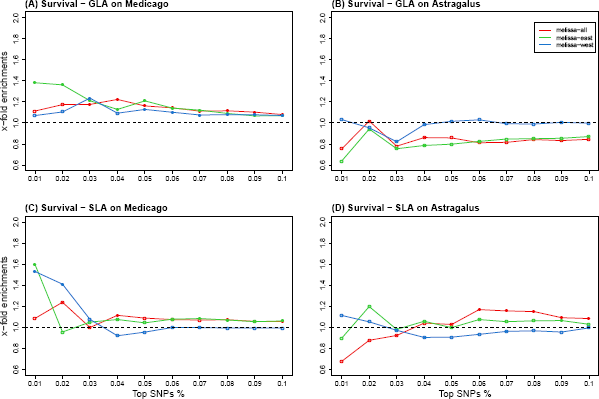
Line plot shows x-fold enrichments across quantiles for overlapping SNPs between survival-associated SNPs in rearing experiment and host-associated SNPs in nature for *ran2B*. Open circles indicate P > 0.05 and filled circles indicate P < 0.05.

**Figure S11:**
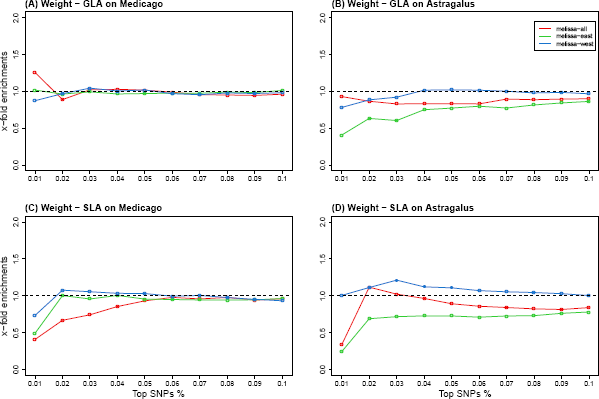
Line plots show x-fold enrichments across quantiles for overlapping SNPs between weight-associated SNPs in the rearing experiment and host-associated SNPs in nature for *ran2B*. Open circles indicate P > 0.05 and filled circles indicate P < 0.05.

**Figure S12:**
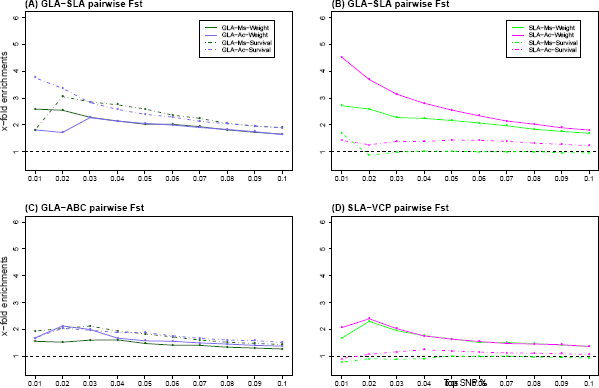
Line plots show x-fold enrichments across quantiles for overlapping SNPs between performance-associated SNPs in the rearing experiment and pairwise Fst-associated SNPs in nature for *ran1*. Open circles indicate P > 0.05 and filled circles indicate P < 0.05.

**Figure S13:**
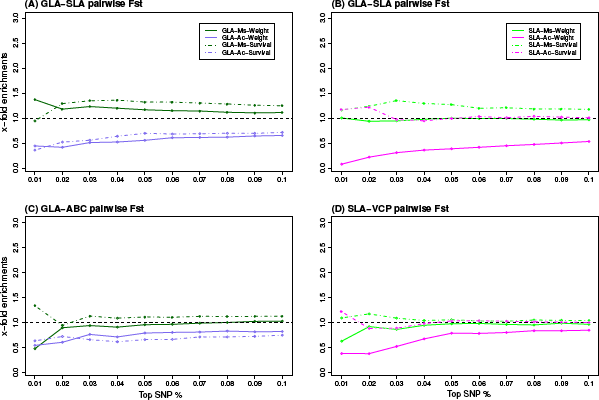
Line plot shows x-fold enrichments across quantiles for concordance in effect signs for overlapping SNPs between performance-associated SNPs in rearing experiment and pairwise Fst-associated SNPs in nature for *ran2A*. Open circles indicate P > 0.05 and filled circles indicate P < 0.05.

**Figure S14:**
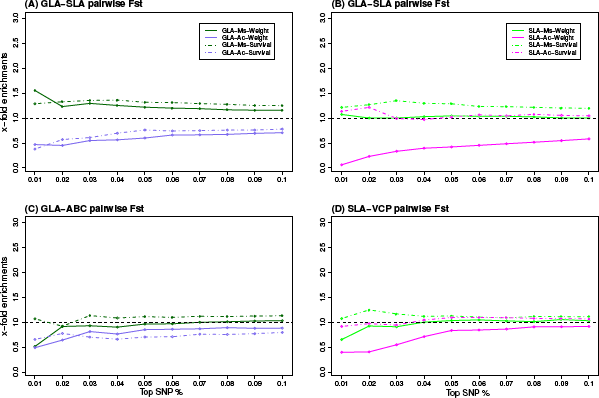
Line plots show x-fold enrichments across quantiles for concordance in effect signs for overlapping SNPs between performance-associated SNPs in the rearing experiment and pairwise Fst-associated SNPs in nature for *ran2B*. Open circles indicate P > 0.05 and filled circles indicate P < 0.05.

